# Simultaneous tracking of division and differentiation from individual hematopoietic stem and progenitor cells reveals within-family homogeneity despite population heterogeneity

**DOI:** 10.1101/586354

**Authors:** Tamar Tak, Giulio Prevedello, Gaël Simon, Noémie Paillon, Ken R. Duffy, Leïla Perié

## Abstract

The advent of high throughput single cell methods such as scRNA-seq has uncovered substantial heterogeneity in the pool of hematopoietic stem and progenitor cells (HSPCs). A significant issue is how to reconcile those findings with the standard model of hematopoietic development, and a fundamental question is how much instruction is inherited by offspring from their ancestors. To address this, we further developed a high-throughput method that enables simultaneously determination of common ancestor, generation, and differentiation status of a large collection of single cells. Data from it revealed that while there is substantial population-level heterogeneity, cells that derived from a common ancestor were highly concordant in their division progression and share similar differentiation outcomes, revealing significant familial effects on both division and differentiation. Although each family diversifies to some extent, the overall collection of cell types observed in a population is largely composed of homogeneous families from heterogeneous ancestors. Heterogeneity between families could be explained, in part, by differences in ancestral expression of cell-surface markers that are used for phenotypic HSPC identification: CD48, SCA-1, c-kit and Flt3. These data call for a revision of the fundamental model of haematopoiesis from a single tree to an ensemble of trees from distinct ancestors where common ancestor effect must be considered. As HSPCs are cultured in the clinic before bone marrow transplantation, our results suggest that the broad range of engraftment and proliferation capacities of HSPCs could be consequences of the heterogeneity in their engrafted families, and altered culture conditions might reduce heterogeneity between families, possibly improving transplantation outcomes.

## Introduction

The hematopoietic system has long since served as a reference model for stem cell biology, with understanding garnered from the study of hematopoietic stem cells (HSCs) successfully transferred to the clinic. In order to maintain blood cell production, rare self-renewing HSCs produce differentiated cells called multi-potent progenitors (MPPs), which proliferate and differentiate through an amplifying cascade of increasingly committed progenitors, ultimately resulting in all mature blood cell types. Underpinning this traditional model, which was mostly uncovered through murine studies, is the assumption that the HSC pool is maintained through a process of asymmetric division that results in one HSC and one MPP, while MPPs form a transient cell type that cannot persist indefinitely and must ultimately differentiate.

Recent studies have challenged this theory in multiple distinct directions. It is well established that HSCs can sequentially reconstitute the blood system of several hosts [1], leading to the inference that HSCs must be able to maintain themselves [2]. When observed with time-lapse imaging, self-renewal has been seen to occur through both symmetric and asymmetric cell division [3–6], which can be influenced by extrinsic signals [5, 6]. Steady state *in situ* lineage tracing studies have also suggested that MPPs are capable of self-renewal [7, 8]. In addition, HSCs have been shown to differentiate without division into megakaryocytes *in vitro* [9], and common myeloid, megakaryocyte and erythroid progenitors *in vivo* [10]. Together, these findings not only questioned the necessity for HSCs to undergo asymmetric division, but they also queried the explicit link between division and differentiation.

Evidence for the multi-potency of HSCs and MPPs has historically derived from *in vitro* colonies assays and transplantation experiments [11–14]. Recent single cell transplantation and cellular barcoding experiments have revealed that only a few HSCs reconstitute all of the blood lineages, with the rest being either restricted in the number of lineages they produce [15–18] or having a bias or imbalance in the proportion of cell types they create [19, 20]. As examples of lineage restriction, it has been reported that some HSCs produce only megakaryocytes [17, 18], while others produce only myeloid cells, megakaryocytes and erythrocytes [15]. Those data suggest that HSCs are a heterogeneous population, where each one of them may be committed to the production of only a few lineages, possibly through lineage priming or externally through instruction from a niche. Similarly, transplanted barcoded MPPs have been reported to produce heterogeneous patterns of restricted cell types [21], suggesting that lineage restriction may occur early in the hematopoietic tree, in the pool of HSCs and MPPs [22].

Altogether, it is presently unclear how symmetric and asymmetric division combines with early lineage commitment to generate down-stream diversity, and a fundamental question is how much instruction is inherited by offspring from an ancestral HSC or MPP. That matter has not been addressed previously due to technical limitations. Tackling it requires an experimental system that enables the simultaneous identification of cells that are descendent from a common ancestor, the number of divisions that has led to each of them, and their differentiation status. Towards that end, we further developed a recently published division-dye multiplex system that was introduced for the study of lymphocytes [23, 24], making it suitable for the study of hematopoietic system.

Amongst other findings, the data from our study revealed that while there is substantial population-level heterogeneity, cells that derived from a common ancestor were highly concordant in their division progression and there were significant familial effects on differentiation. This similarity is primarily propagated through divisions resulting in siblings of the same cell type, although a small number of asymmetries are sufficient to break perfect symmetry, so that individual ancestors create a diversity of lineages. Our data establishes that early lineage commitment can be inherited from individual HSCs and MPPs, and that the resulting diversity of lineages is produced by a heterogeneous collection of cell families that are individually homogeneous. While the current single cell revolution has led to significant breakthroughs in understanding stem cell biology, our data suggest that common ancestor effects are significant and must also be considered.

## Results

Defining a family as all descendants from a marked ancestor cell, the experimental method employed here is based on the observation that by labeling cells with distinct combinations of division diluting dyes, such as 5-(and 6)-carboxyfluorescein diacetate succinimidyl ester (CFSE) and CellTrace Violet (CTV), through the use of flow cytometry it is possible to simultaneously determine each cell’s family membership, generation (i.e. number of cell divisions), and phenotype [23, 24] (Figure 1). In the present application, by index sorting labeled cells, we could also relate down-stream familial fate to ancestral cell surface expression. We chose to investigate familial division and differentiation in early hematopoietic differentiation from HSCs, focusing on three phenotypically defined cell populations: c-Kit^+^Sca-1^+^SLAM^+^Flt3^−^ (SLAM-HSC), c-Kit^+^Sca-1^+^SLAM^−^Flt3^−^ (ST-HSC) and c-Kit^+^ Sca-1^+^SLAM^−^Flt3^+^ (MPP) (Figure 1).

**Figure 1.**
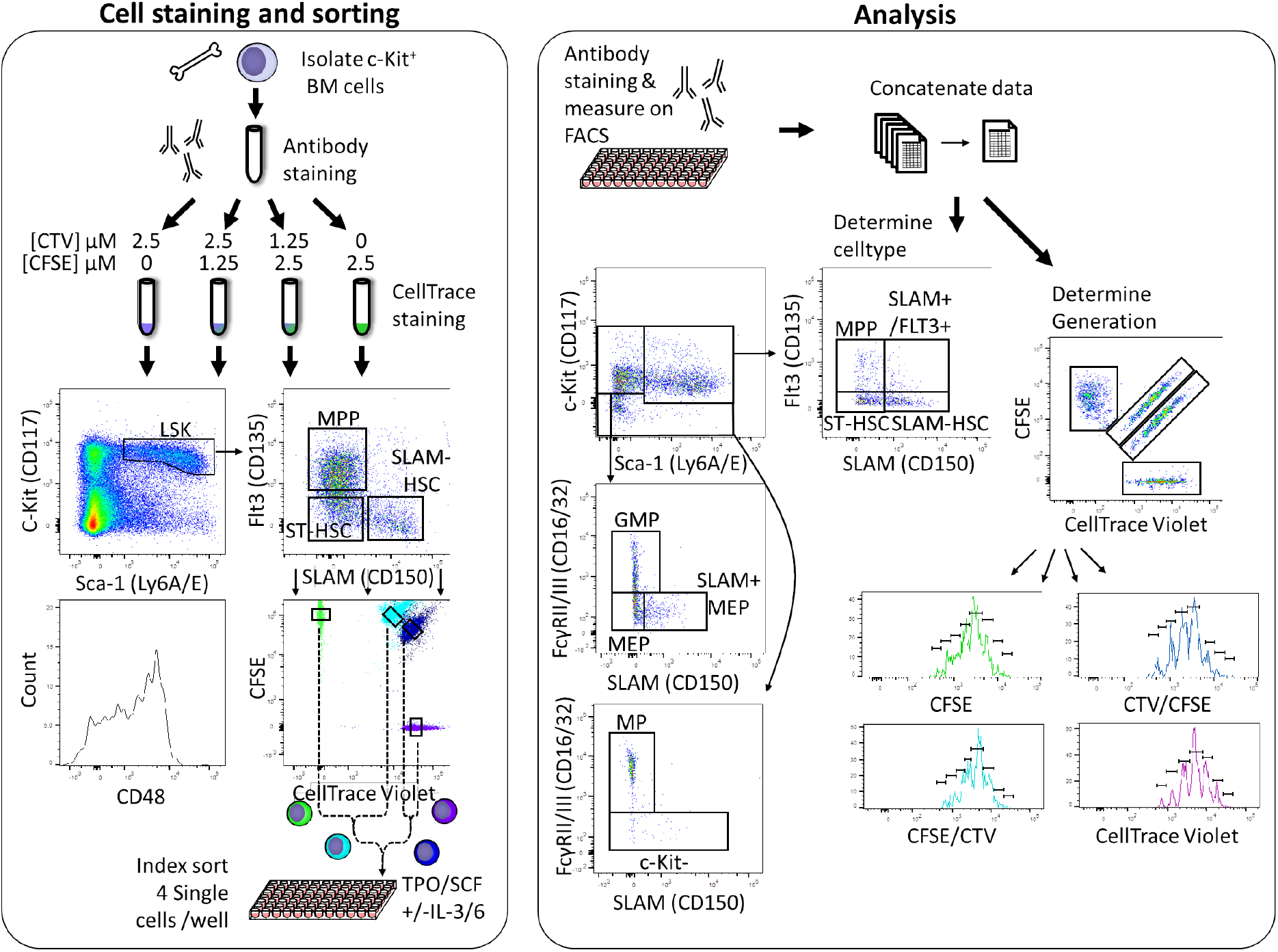
High throughput simultaneous division and differentiation tracking per-ancestor. A single cell-suspension was obtained by flushing femurs, tibia and iliac crests. Cells were stained with fluorescently labeled antibodies for phenotypic identification. The cell suspension was then split into four equal parts, each of which was stained with a distinct CFSE and CTV combination. From each of these preparations, a single cell was index sorted into 90 wells of a 96-well plate, resulting in four distinctly colored cells per well. In addition, for each ancestor type, 100 cells of each color combination were sorted into a single well. After 24 or 48h of culture, cells were stained with fluorescently labeled antibodies for phenotypic identification and analyzed on a flow cytometer. The data from all wells were combined and used to set gates for determination of generation number and phenotypic cell type. Those gates were then applied to the data from each well to obtain lineage, division, and differentiation information for each family originating from a founding ancestor cell.

We isolated bone marrow cells, labeled them with four distinguishable combinations of CFSE and CTV, and used fluorescent antibody staining of cell surface markers to determine their phenotype (Figure 1). Wells in a 96 well plate were then seeded with four cells of a single ancestral type (SLAM-HSC, ST-HSC or MPP), one from each of the four CFSE/CTV combinations, and incubated in one of two classic cytokine cocktails (SCF and TPO, +/− IL3 and IL6). At 24 or 48h, cells were harvested from each well and stained with fluorescent antibodies to determine their phenotypic cell type based on the expression of SLAM, Flt3, Sca-1, c-Kit and CD16/32 (Figure 1). By examining each cell’s CFSE and CTV profile, its ancestral cell and generation number was determined (Figure 1 and Methods). To capture early division and differentiation as well as later developments, for each ancestor type (SLAM-HSC, ST-HSC, MPP) 360 initial cells were sorted for analysis at 24h, and 240 initial cells for analysis at 48h (Figure 2A). From the 600 seeded SLAM-HSCs, in total we recovered 358 families (71%) constituting 648 cells, while 343 ST-HSC families (69%) were recovered with 592 cells, and 246 MPP families (49%) with 362 cells (Figure 2A). Over all conditions, 27 families (2.8% of recovered families) had cell numbers that could not have originated from a single ancestor, and so were excluded from analysis.

**Figure 2.**
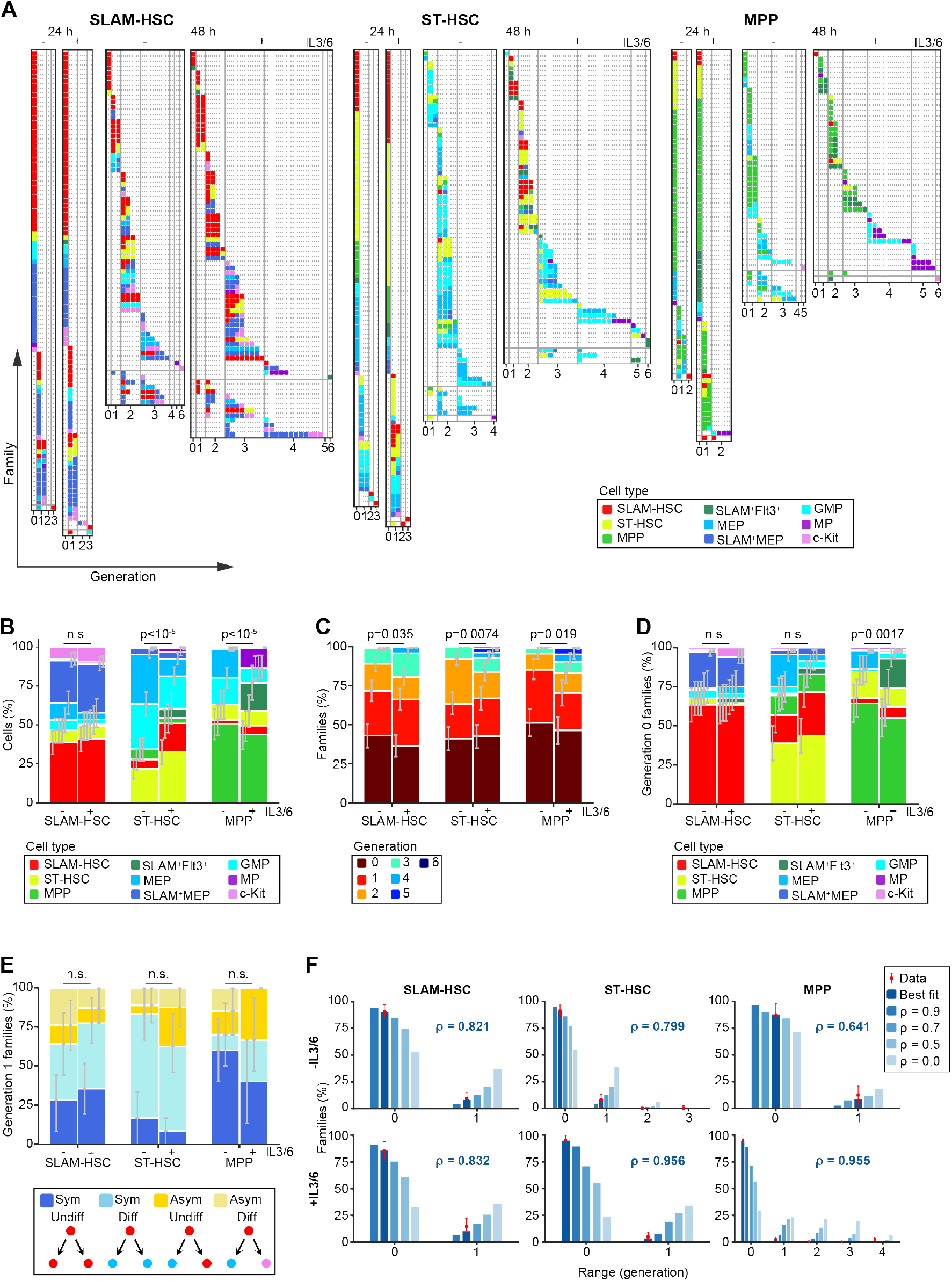
Despite population level heterogeneity, individual HSPC families are substantially homogeneous. Plots are fractionated by each ancestor type (SLAM-HSCs, ST-HSCs, and MPPs) and cocktail (with and without IL3+IL6, indicated by + and − respectively). For (B-F), error bars indicate 95% confidence intervals calculated via basic bootstrap (Methods). Sample sizes and p-values (from permutation tests, see Methods) for each panels can be found in Supplementary Table 1 and 2. (A) Simultaneous visualization of family membership, generation number, and cell type of offspring from initially seeded ancestors harvested at two time points (24 and 48h). Each row presents the offspring from a single ancestor. Columns identify the generation number of each recovered cell, with their phenotypic cell type indicated by color-coding. Rows are sorted in increasing order of the difference between maximum and minimum generations in each ancestor’s family (generational range). (B) Percentage of each recovered cell type. (C) Distribution of the maximum generation per family, as indicated by color-coding. (D) Proportions of recovered cell types for ancestors that have not yet divided. (E) For ancestors that have divided only once and for whom two offspring are recovered, percentage undergoing symmetric and asymmetric division with and without differentiation. (F) Percentage of families with each generational range. The 48h data (red dots) is compared to a mathematical model parameterized by a single coefficient, ρ, which encodes the correlation in whether cells in the same generation within a family divide or cease to divide (Methods). Shown is prediction for both the maximum likelihood best-fit value of ρ and, as a reference, a range of other values of ρ.

At the population level, some offspring underwent no differentiation from their ancestor type while others differentiated. In both culture conditions and for each ancestor type, we obtained a diversity of myeloid cell types, ranging from the initial ancestor to c-Kit^−^ differentiated cells, included c-Kit^+^Sca-1^−^CD16/32^+^ (GMP) and c-Kit^+^ Sca-1^−^SLAM^−^CD16/32^−^ (MEP) cell types (Figure 2A and B). We also detected c-Kit^+^Sca-1^−^CD16/32^−^SLAM^+^ (SLAM^+^MEP) cells, previously described as megakaryocyte and erythroid progenitors [25]. Although all hematopoietic cells have been reported to go through a Flt3 expressing stage [26], we found that no offspring of SLAM-HSCs, and those of very few ST-HSCs, differentiated into Flt3 expressing MPPs in our culture conditions (Figure 2B). Of the seeded ST-HSCs and MPPs, a small number of their offspring were observed to dedifferentiate as they acquire the upstream SLAM-HSC or ST-HSC phenotypes (Figure 2B). As cells at the edge of FACS gates can be incorrectly categorized, we compared the index sorting surface marker expression of cells whose offspring dedifferentiating with those that did not dedifferentiate, but no significant difference was identified (Figure S1A). Thus, these data indicated that phenotypically defined ST-HSCs are capable of dedifferentiating into SLAM-HSCs, and MPPs are capable of dedifferentiating into both ST-HSCs and SLAM-HSCs in our culture conditions. The addition of IL-3 and IL-6 had no impact on the pattern of cell types produced by SLAM-HSCs (Figure 2B). It did, however, change the pattern of cell types produced by ST-HSCs and MPPs, with an increase in ST-HSC dedifferentiation to SLAM-HSC, and an increase in MPP differentiation into SLAM^+^Flt3^+^ cells, indicating that SLAM^+^Flt3^+^ cells can arise by gain of SLAM expression.

At the level of individual families, we observed substantial heterogeneity in division history (Figure 2C), from families that did not proliferate to those with cells that had undergone 6 divisions. Consistent with earlier observations [27], after 24h most cells either remained undivided or were in generation one, with a few cells having undergone two or more divisions (Figure 2A). At 48h over 90% of the cells had undergone at least one division. For all three sorted ancestor cell types, we found that addition of IL-3 and IL-6 led to a statistically significant increase in proliferation (Figure 2C), as previously described [28, 29]. Exploring the relationship between division and differentiation, in the culture without IL3 and IL6, the proportion of undivided ancestors that differentiated were 36.5% from SLAM-HSCs, 61.1% from ST-HSCs, and 35.6% from MPPs (Figure 2D), which was in agreement with previous reports [9, 10]. The addition of IL3 and IL6 didn’t drastically change those values: 36.8% from SLAM-HSCs, 56.3% from ST-HSCs, and 44.8% from MPPs. SLAM-HSCs preferentially differentiated without dividing into SLAM^+^MEP, as reported previously [10]. On comparing the surface marker expression of differentiated and not-differentiated non-divided cells (Figure S1B), Sca-1^high^ SLAM-HSCs were more likely not to differentiate, in agreement with a previous report [30]. The addition of IL-3 and IL-6 only significantly impacted the differentiation pattern of the progeny of MPPs. These results show that families are heterogeneous in their division pattern, and that a non-negligible fraction of ancestors differentiates without dividing.

A desirable feature of our experimental system is that it is can capture a large number of siblings after a single division, enabling the quantification of symmetric versus asymmetric division. In our system, we defined four distinct types of symmetric or asymmetric division depending on whether the offspring included a differentiated cell (Figure 2E). Symmetric undifferentiated division produces two cells of the ancestor type. Asymmetric undifferentiated division, which would be the classically defined asymmetric division in the stem cell community, produces one cell of the ancestor type and one differentiated cell. Similarly, symmetric differentiated division produces two cells of the same differentiated type, and asymmetric differentiated division produces two cells of the distinct differentiated type. Note that asymmetric division cannot be distinguished from symmetric division followed by differentiation without division of one of the daughter cells. Pooling data over ancestor types, 70.7% of first divisions were symmetric, with MPPs performing mostly symmetric undifferentiated divisions (51.4%), and ST-HSC performing mostly symmetric differentiated divisions (59.5%). SLAM-HSCs self-renewed primarily through symmetric undifferentiated division (32.1%), but also through asymmetric undifferentiated divisions (10.7%). Asymmetric divisions occurred for 28.6% of SLAM-HSCs, 28.6% of ST-HSCs and 31.4% of MPPs, consistent with previous reports [31]. No statistical difference was found between ancestors cultured with or without IL3 and IL6. The pattern of cell types produced after one division (Figure S2) was similar to the pattern including all divisions (Figure 2B), suggesting that the diversity of cell types can be produced by a heterogeneous collection of cell families through symmetric divisions.

To investigate familial effects on division progression, we examined the generation numbers of cells derived from single ancestors. We found that families are highly concordant, with 81% of the 223 families that divided more than once having all of their cells in the same generation (range 0, Figure 2F), and only four of those families (1.8%) containing cells that were more than one generation apart (range>1, Figure 2F). As not all cells are necessarily recovered from wells and sampling effects could potentially make families look more concordant, by fitting a mathematical model to the 48h data that takes the empirically determined sampling into account we performed a quantitative assessment of the correlation in division progression decisions necessary by cells within a family to explain the data. In the model, the probability that cells within a family divide or stop dividing at each generation is coupled by a single correlation parameter ρ (Methods). The marginal probability of division was set to mimic the observed proportion of division progressing cells, and the correlation parameter, ρ, was then fit to per-ancestor data (Methods). High correlation coefficients of between 70-90% (Figure 2F) resulted in the best fit to the measured familial ranges, with the exception of MPPs cultured in medium without IL-3 and IL-6 for which no reliable estimate could be made. For guidance, the pattern of range values from the model for different correlation coefficients are also illustrated in the figure. Thus, this analysis establishes that division progression is highly concordant within SLAM-HSC, ST-HSC and MPP families, while being heterogeneous between them.

Although differentiation without division was observed, in general differentiation progressed in tandem with division (Figure 3A). To visualize changes in cell surface marker expression, we used the Uniform Manifold Approximation and Projection algorithm (UMAP) [32] on the combined surface marker expressions of all cells obtained at both 24 and 48h (Figure 3B and Figure S3). When we mapped cell types determined by traditional gating onto the UMAP (Figure 3C), a smooth transition was observed from SLAM-HSCs at the bottom, with further differentiated cells towards the top. GMPs and c-Kit^−^Sca-1^−^CD16/32^+^ (Myeloid Progenitors, MPs) appear on the top left, and MEP and SLAM^+^-MEP on the top right. More numerous GMPs, MPs and MEPs were seen at the top of the UMAP at 48h than 24h, suggesting that it takes between 24 to 48h to fully differentiate into Sca1^−^ progenitors. In addition, all three ancestral cell types remain present at 48h, indicating, in particular, that HSCs can remain in an undifferentiated state for the duration of the experiment, even if their offspring experience three rounds of division. On plotting the generation numbers of offspring from each ancestor cell type on the UMAP (Figure 3D and Figure S4), SLAM-HSCs appeared to be primed toward the production of MEPs and SLAM^+^MEPs while still generating some ST-HSCs, whereas MPPs were more primed toward GMPs, and ST-HSCs showed a more even distribution between the two lineages. Differentiation without proliferation appeared as dark red dots outside of the regions of the sorted ancestor cells, and self-renewal divisions as red, orange and blue dots in the region of ancestor cells. SLAM^+^MEPs were observed to be generated without division, as well as in one to three divisions from SLAM-HSCs. This path arose more rapidly in our data than previously reported [33].

**Figure 3.**
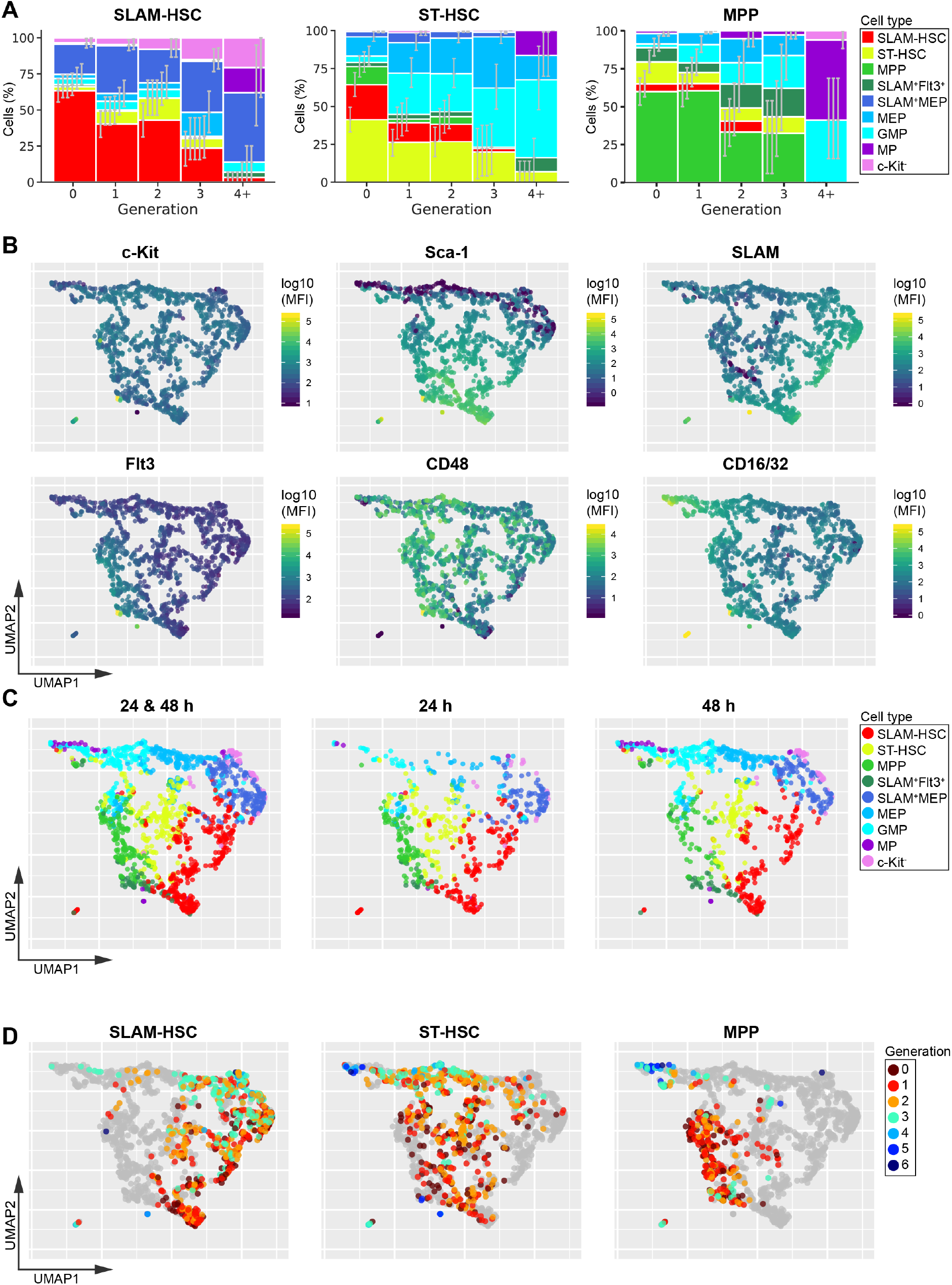
Differentiation and division progress in tandem. (A) Percentage of cells at each differentiation stage for each generation for each ancestor type. Error bars indicate basic bootstrap 95% confidence intervals (Methods). Sample sizes for this panel can be found in Supplementary Table 3. (B) The Uniform Manifold Approximation and Projection (UMAP) algorithm was applied to the data pooled from all time-points, conditions and ancestor cell types. Each cell is projected into the UMAP coordinates and color-coded according to the log of their median fluorescence intensity for c-Kit, Sca-1, SLAM, Flt3, CD48 and CD16/32 at the time of analysis (see also Figure S3, and for each ancestor cell type plotted separately on the UMAP see Figure S5). (C) Projection of traditionally gated data onto the UMAP, pooled data (left), 24h (middle) and 48h (right). (D) Projection of cell generation number data onto the UMAP for each ancestor type (see also Figure S4). Sample sizes for panels B-D can be found in Supplementary Table 1.

Descendants from a common ancestor were not only highly concordant in their generation numbers, but they also exhibited significant similarity in differentiation outcome. At 24h, most families were composed of only one cell type (Figure 4A), but at 48h more families produced several cell types (Figure 4B) indicating that downstream asymmetries in the fate occurs after a largely symmetric first division (Figure 2E). Permutation tests on phenotypically defined cell types revealed that families exhibited significant more similarity than would be expected if there was no family component, both at 24h (Figure 4A) and at 48h (Figure 4B). To visualize the ancestral impact on differentiation, on the UMAP we displayed the offspring from the 15 families with the highest number of recovered cells (Figure 4C). Thus, SLAM-HSC, ST-HSC, and MPP families are highly concordant in division and share similar differentiation outcomes *in vitro*, while population level diversity in proliferation and cell types arises from heterogeneity across families.

**Figure 4.**
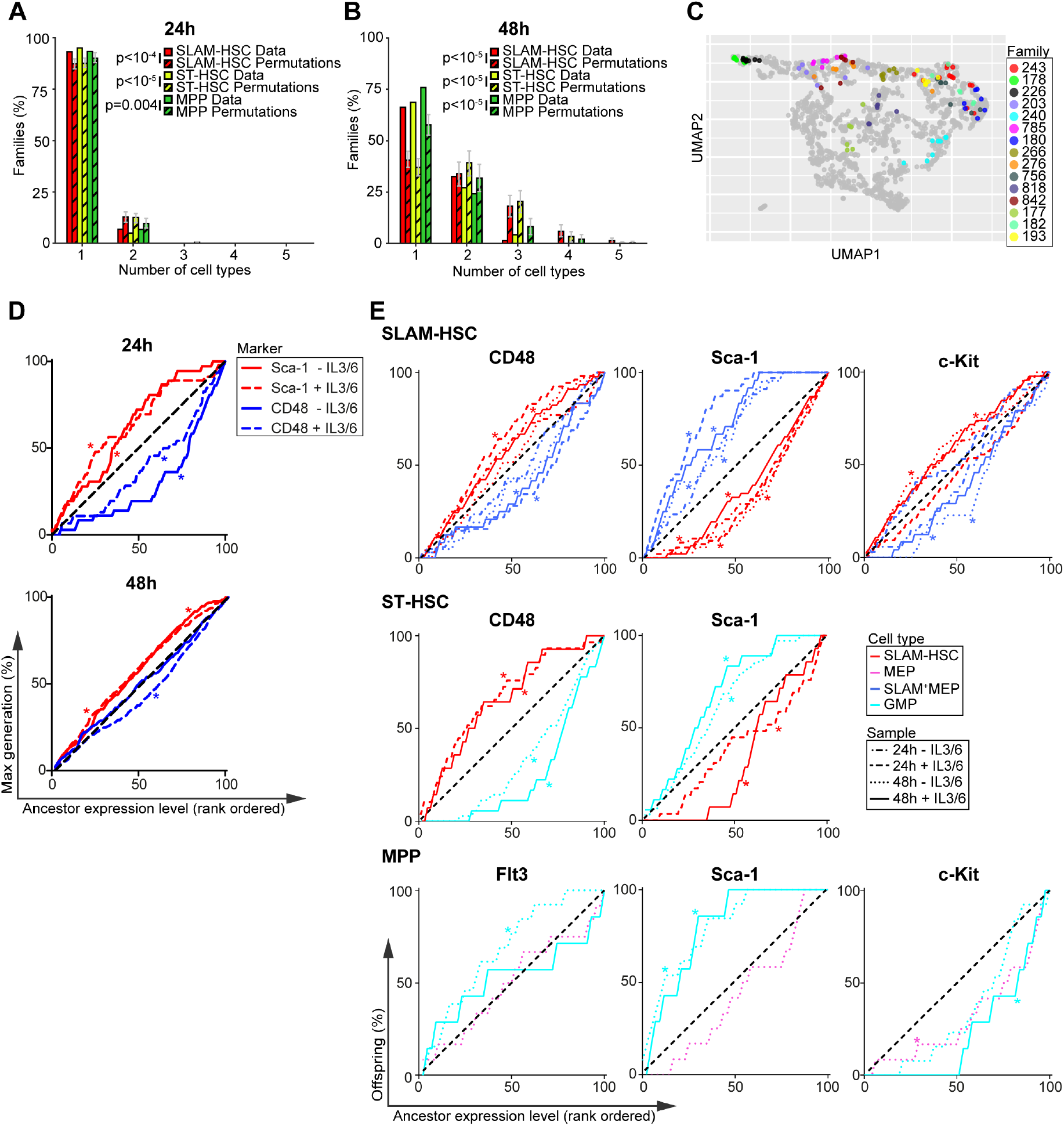
Families are highly concordant in differentiation. (A-B) Number of cell types per family in the observed data (bars with no pattern) compared with the average of 250,000 permutations of the data (bars with pattern) at 24h (A) and 48h (B). Error bars indicate 95% confidence intervals based on permutations (see Methods). (C) Cells from the 15 families with the largest number of cells are color-coded by family and projected onto the UMAP in Figure 3. (D) The cumulative percentage of the maximum division of offspring from ancestor SLAM-HSCs rank-ordered by their expression of CD48 (blue) or Sca-1 (red) during sort. (E) The cumulative percentage of offspring, presenting a given cell type, from ancestor cells rank-ordered by increasing cell surface marker expression. * indicates a significant deviation from the diagonal (black) as determined by Jonckheere’s Trend Test (p-values in Supplementary Table 5 and 6). Sample sizes for all panels can be found in Supplementary Table 1.

The within-family homogeneity in division and differentiation could be intrinsically present in ancestor cells or extrinsically instructed by cytokines in the cocktail. Comparison of the median fluorescent intensities of markers from ancestors during sort with those obtained from daughters at each of the two time points (Figure S6A) revealed clear correlation between the two, supporting the hypothesis that ancestor surface expression markers were instructive in the within-family homogeneity in division progression and differentiation. To further investigate that hypothesis, we first explored the relationship between cell surface expression on the ancestor and division progression. Rank ordering ancestors from the least to greatest expression level for a given marker (Figure 4D, Figure S6B), the cumulative sum of the maximum division of their offspring would be expected to fall near the diagonal if there were no relationship between an ancestral expression level and division progression. If there was a negative relationship, where low expression of a given marker on the ancestor corresponded to more division progression, the cumulative sum of the maximum division would be expected to initially overshoot above the diagonal. In contrast, if there was a positive relationship, instead there would be an initial undershoot of the diagonal. The statistical significance of divergence from the diagonal was tested using Jonckheere’s Trend Test [34] (Supplementary Table 5), which challenges a null hypothesis of no relationship in median against alternative hypotheses of negative or positive trend. Across both cocktails and time-points, only cell surface markers on ancestral SLAM-HSCs were consistently instructive for division progression. CD48 correlated positively, with its strongest effect at 24hr, and Sca-1 correlated negatively, while at 48h c-Kit correlated positively (Figure 4D and Figure S6B). Notably, for both ST-HSCs and MPPs, even though the family data clearly indicate that there is a familial component to division progression (Figure 2F), none of the phenotypic markers exhibited strong correlation (Figure S6B), indicating the need to identify other factors.

We then explored the relationship between ancestral marker expression and familial differentiation (Figure 4E, Figure S7 and Supplementary Table 6). For SLAM-HSCs, at 24 and 48h in both cocktails, Sca-1 expression provided a strong positive correlation to selfrenewal and a negative one to production of SLAM^+^MEPs (Figure 4E). At 48h, c-Kit presented the inverse dependency to Sca-1. At 24 and 48h, CD48 correlated negatively to self-renewal and positively to production of SLAM^+^MEPs when IL-3 and IL-6 is added. As cocktail composition did not have a major impact on the relationship between familial fate and ancestral expression of Sca-1 and c-Kit, these results were suggestive that c-Kit and Sca-1 expression levels of SLAM-HSCs act as intrinsic markers for both familial progression and differentiation with high Sca-1 expression and low c-Kit expression leading to less division [35–37] and less differentiation [38], and potentially resulting in better engraftment [35, 36, 38]. While low ancestral CD48 expression level has been reported to result in less division [39, 40], our data indicates its relationship to differentiation is dependent on extrinsic signals.

For ST-HSCs and MPPs we found little evidence of correlation of ancestral expression to division progression or self-renewal, but the same was not true of differentiation. For ST-HSCs, the ancestral level of CD48 and Sca-1 consistently correlated negatively and positively, respectively, with de-differentiation to SLAM-HSC in the cocktail with IL-3 and IL-6 at both time points (Figure 4E and S7, and Supplementary Table 6). Differentiation to GMP, which occurred only in 48h data, correlated positively and negatively with the ancestral level of CD48 and Sca-1 respectively (Figure 4E) [37]. Therefore, differentiation to GMP from ST-HSC was dependent on the parental level of CD48 and Sca-1, whereas the dedifferentiation to SLAM-HSC is dependent on both extrinsic factors (IL-3 and IL-6) and the intrinsic ancestral level of CD48 and Sca-1. The differentiation from MPPs to GMPs that was observed to occur by 48h correlated negatively with Sca-1 ancestral expression [37] in both cocktails. It also negatively correlated with Flt3 ancestral expression, but only in the cocktail without IL-3 and IL-6 (Figure 4E). In the presence of IL-3 and IL-6, instead, differentiation from MPP to GMP positively correlated with c-Kit ancestral expression. Differentiation from MPP to MEP occurred only at 48h in the cocktail without IL-3 and IL-6, and then correlated positively with c-Kit (Figure 4E). Thus differentiation to GMP from MPP is dependent on the intrinsic ancestral level of Sca-1, whereas the differentiation to MEP is dependent on both extrinsic factors (IL-3 and IL-6) and the intrinsic ancestral level of c-Kit. Overall, the concordance in division and similarity in fate within families is partially explained by the surface expression marker used to phenotype ancestors, but both intrinsic and extrinsic factors act to direct familial fate.

## Discussion

We further developed a recently published high-throughput method that enables simultaneously determination of common ancestor, generation and differentiation status of a large collection of single cells. Its use with HSPCs led us to the new discovery that, despite substantial population-level heterogeneity amongst offspring, cells derived from a single ancestor are highly concordant in their division progression and exhibit familial effects on differentiation. The restriction in differentiated cell types within each family is propagated through mostly symmetric first divisions followed by a few downstream asymmetries in fate. Although each family diversifies to some extent, the overall collection of cell types observed in a population is largely composed of homogeneous families from heterogeneous ancestors. This finding opens new avenues and challenges for the hematopoietic field. The generation of a diversity of cell types is presently assumed to result from a diversification within a family, and methods for inferring differentiation trajectories using single cell RNA sequencing data from snapshot data assume that cells all behave independently [41, 42]. Consistent with previous observations of early lineages decisions [22, 43, 44], our findings establish that familial dependencies that are currently unmeasured exist within the population, and call for a revision of that assumption and subsequent analysis. Ancestral cell surface expression of markers used for phenotyping serve as correlates that partially predict some of these familial properties, but, in particular, a correlate that explains the highly heritable division progression of ST-HSC and MPP families is not contained within them. It is also the case that extrinsic properties such as cytokine signaling can play an instructive role, altering and reshaping the observed familial effects.

As in the clinic HSPCs are cultured before bone marrow transplantation, our results indicate that the broad range of engraftment and proliferation capacities of HSPCs could be consequences of the heterogeneity in their engrafted families. That suggests that altered culture conditions might reduce heterogeneity between families and possibly improve transplantation outcomes. Indeed, changing the composition of the population of committed HSPC might be a mechanism to directly alter the balance of lineage production, with therapeutic applications that could benefit the treatment of both leukemia and genetic immune disorders.

## Methods

### Mice and cell isolation

All the experimental procedures were approved by the local ethics committee CEEA-IC (Comité d’Ethique en expérimentation animale de l’Institut Curie) under approval number DAP 2016 006. BM cells were obtained from wild-type C57BL/6 of 8-16 weeks of age by bone flushing of femur tibia and iliac crest. Bone marrow cells were MACS enriched for c-Kit^+^ cells using CD117 MicroBeads Ultrapure (Miltenyi Biotec cat #130-091-224) according to manufacturer’s protocol.

### Division tracking and surface marker labeling of HSPC

c-Kit enriched BM cells were stained with CD135 (Flt3) PE (eBiosciences 12-1351-82), Sca-1 PE-CF594 (BD Biosciences, 562730), CD117 (c-Kit) APC (Biolegend 105812), CD150 (SLAM) PC7 (Biolegend 115914) and CD48 APC-Cy7 (Biolegend 103432) for at least 40min at a concentration of 1 to 100. Fluochromes were chosen for having very little spillover from the bright CellTrace dyes. Subsequently, cells were split into 4 equal fractions and simultaneously stained in PBS at 37°C for 15 min with either 2.5 μM CellTrace CFSE (ThermoFisher Scientific C34554), 2.5 μM CellTrace Violet (CTV) (ThermoFisher Scientific C34557), 2.5 μM CFSE together with 1.25 μM CTV or 2.5 μM CTV together with 1.25 μM CFSE (see Fig. 1A) as adapted from [23]. The reaction was quenched by washing with 2×5mL ice cold RPMI supplemented with 10% FCS.

c-Kit^+^ Sca-1^+^SLAM^+^Flt3^−^ (SLAM-HSC), c-Kit^+^Sca-1^+^SLAM^−^Flt3^−^ (ST-HSC) and c-Kit^+^Sca-1^+^SLAM^−^/Flt3^+^ (MPP) were sorted directly into U-bottom 96-well plates containing cell culture media using an Aria III cell sorter (BD Biosciences). For each cell type we sorted 4 single cells, one for each of the CellTrace stain combinations, into each well. In total 30 wells (120 single cells) were sorted per cell type per plate, with 3 replicates for analysis at 24h and 2 replicates for analysis at 48h. In addition, we sorted 100 cells of each cell type into one well for both culture conditions, as well as 1000 c-Kit^+^ cells into another well. During the sort of single cells, fluorescence intensities of each surface marker were recorded using the index sorting function.

### In vitro cell culture

Cells were cultured at 37°C under 5% CO2 in 100 μl of StemSpan serum-free expansion medium (STEMCELL technologies 9650) supplemented with cytokine cocktails aimed at proliferation (50 ng/ml murine recombinant thrombopoietin (TPO, Sigma-Aldrich SRP3236-10UG) and 100ng/ml Stem Cell Factor (SCF)) or proliferation and differentiation (50 ng/ml TPO, 100 ng/ml SCF, 20 ng/ml IL-3 and 100 ng/ml IL-6) [5, 28].

### Division and expression marker analysis of cell progeny

After 24 or 48h of incubation, cells were stained for 40 min in RPMI 1640 supplemented with 10% FBS using the same antibody combination as for sorting, with several modifications: CD48 BUV395 (BD biosciences 740236) was used instead of CD48 APC-Cy7, Sca-1 APC-Cy7 (Biolegend 108125) instead of PE-CF594 and CD16/32 BV711 (BD biosciences 101337) was added to the cocktail at a concentration of 1 to 100. To minimize loss of cells, only two washing steps were performed. Cells were analysed at 4°C using a ZE5 Flow cytometer (BioRad) set to sample wells until running dry, with a recovery estimate of circa 70% per well (beads-based estimate, data not shown).

### Cell type and generation assignment

For data analysis of FACS data we pooled all the data from a single experiment using the concatenate function in FlowJo (FlowJo, LLC version 10.4.2).

For cell type assignment, gates were set on concatenated data and then applied to clonal data as shown in Figure 1. In short, cells were separated from debris by their FSC/SSC profile. Subsequently, cells were identified by gating on CD16/32, c-Kit, Sca1, Flt3 and SLAM expression as c-Kit^+^Sca-1^+^SLAM^−^Flt3^−^ (SLAM-HSC), c-Kit^+^Sca-1^+^SLAM^+^Flt3^−^ (ST-HSC), c-Kit^+^Sca-1^+^SLAM^−^Flt3^+^ (MPP), c-Kit^+^Sca1^+^SLAM^+^Flt3^−^ cells (SLAM+/Flt3+) c-Kit^+^Sca-1^−^CD16/32^+^ Granulocyte-Macrophage Progenitors (GMP), further differentiated c-Kit^−^ Sca1^−^CD16/32^+^ Myeloid Progenitors (MP) or c-Kit^−^CD16/32^−^ progenitors (c-Kit^−^), c-Kit^+^Sca-1^−^SLAM^−^CD16/32^−^ Megakaryocyte-Erythroid Progenitors (MEPs) and lastly c-Kit^+^Sca-1^−^SLAM^+^CD16/32^−^/ (SLAM^+^-MEPs), which have been shown to contain both pre-CFU-E and pre-Megakaryocyte/Erythrocyte progenitors.

To determine the generation (i.e. the number of divisions since labeling) of cells, cells were first divided into their four distinct CFSE/CTV combinations. For cells stained with either CFSE or CTV alone, generation gates were determined based on histograms of their dye florescence on a logarithmic Scale in FlowJo (FlowJo, LLC version 10.4.2). For cells stained with a both CFSE and CTV, we rotated the CTV/CFSE coordinates, on a logarithmic Scale, by 45 degrees anticlockwise so that division dilution proceeded in parallel to the horizontal. That is, with *x* and *y* denoting the coordinates of CTV and CFSE levels, the histogram was calculated over a new x-axis coordinate

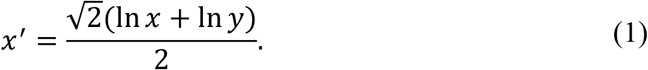

Generation gating was then determined based on the florescence histogram on the new x’-axis on the merged data of wells from the same experiment.

### Data visualisation by UMAP

Before processing, flow cytometric fluorescence intensity values from all experiments were pooled and transformed by an arcsinh(x/100) transformation. On these data we performed Uniform Manifold Approximation and Projection (UMAP) [45] using the R implementation in the UMAP package (version 0.2.0.0) with default parameters. The output of the UMAP was visualized using the ggplot 2 package (version 3.0.0) in R (version 3.4.3).

### Statistical testing by permutation

Using the data underlying Figures 2B-E, we challenged the hypothesis that division and differentiation are independent of culture condition (i.e. with or without IL-3 and IL-6) for cells derived from the same ancestor type (SLAM-HSC, ST-HSC or MPP). To that end, we adapted the permutation test [46] framework proposed in [24].

To challenge the null hypotheses that differentiation was independent of the culture condition, using the data underlying Figure 2B we compared the population proportion per cell type. For notational purposes, the data were represented as a sequence 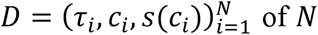 of *N* cells, where the *i^th^* cell was identified by: cell type *τ_i_*; clone *c_i_*; and culture condition of the clone *s*(*c_i_*). To assess the independence of cell types *J* = {*τ_i_,i* = 1,…,*N*} from partition labels *l* ∈ {1,…,*L*} (relative to culture condition), the statistic *T* of the data *D* was defined as the log-likelihood statistic of the G-test for the contingency table *O*, such that 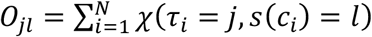 with *χ*(*A*) = 1 if the event *A* holds true and 0 otherwise. The G-test statistic is classically used for the testing of independence between two sets of categories (*J* and {1,…,*L*}) partitioning the data counts [46]. Therefore,

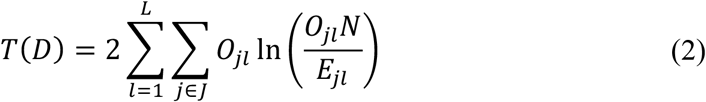

where 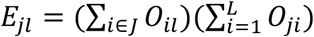.

Under the null hypothesis that differentiation was not impacted by culture condition, *D* is equally likely as a dataset 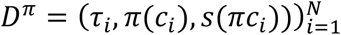 transformed by the action of any permutation *π* ∈ *Q* of the set of clonal labels {*c_i_, i* = 1,…,*N*}. As a consequence, using Monte Carlo approximation we estimated the p-value for the right-tailed test as

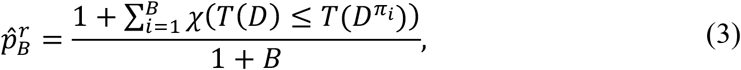

where *B* = 250,000 and *π*_1_,…,*π_B_* were uniformly and independently sampled from *Q*.

To challenge the null hypotheses that clonal division was independent of the culture condition, for the data underlying Figure 2C we compared the distribution of the maximum generation reached by each clone. For these procedures, it sufficed to follow the same rationale as for the tests related to Figure 2B, but for the dataset 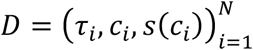 of *N* clones, where *τ_i_* is the maximum generation of the *i*^th^ clone. In particular, the testing statistic *T* was defined as in (2) and the subsequent p-value was estimated as in (3).

To challenge the null hypotheses that differentiation without division was independent of the culture condition for the data underlying Figure 2D, we compared the proportions of cell types of undivided cells (i.e. those in generation 0). For these procedures, it sufficed to follow the same rationale as for the tests related to Figure 2B and 2C, with 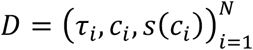 the sequence of *N* clones in generation 0, where *τ_i_* identifies the type of the unique cell in clone *c_i_*. In particular, the testing statistic *T* was defined as in (2) and the subsequent p-value was estimated as in (3).

To challenge the null hypotheses that the pattern of first division was independent of the culture condition, for the data underlying Figure 2E we compared the proportion of division types among clones recovered with two cells in generation 1. For these procedures, it sufficed to follow the same rationale as for the tests in Figure 2B-D, with 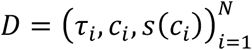 as the dataset of *N* clones with two cells generation 1, where *τ_i_* records the pattern of division of the clone *c_i_* as one out of four possibilities (outlined in Figure 2E). The test statistic *T* was defined as in (2) and the subsequent p-value was estimated as in (3).

For the data in a given time point (24 or 48h) underlying Figure 4B, we investigated the clonal effect on differentiation by challenging the null hypotheses that differentiation diversity among families from the same ancestor type was independent of clonal membership. In particular, as the cells from the data were found in different generations, we sought to take into account that division may have had an impact on differentiation (Figure 3D). These data were identified by the sequence 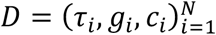 of the *N* cells from the same progenitor, with *τ_i_, g_i_, c_i_* recording the type, the generation and the clonal label, respectively, of the *i*^th^ cell. To test the null hypothesis by permutation, the set of invariant transformations *Q* for *D* should permute, across clones, only cells that were found in the same generation. To this end, *Q* was generated by the functions *π_g_* for *g* ∈ *G* = {*g_i_, i* = 1,…,*N*], such that

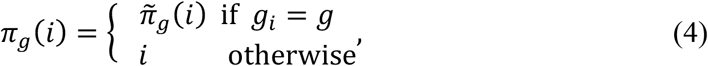

where 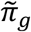 is any permutation of the set {*i* = 1,…,*N: g_i_* = *g*}. Then 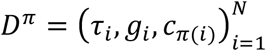. To measure clonal differentiation diversity, we defined the statistic *T* for the average number of cell types per family, that is

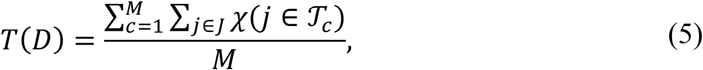

where {1,…,*M*} is the set of all clonal labels, *J* = {*τ_i_: i* = 1,…,*N*} is the set of all cell types observed, and *T_c_* = {*τ_i_, i* = 1,…,*N*: *c_i_* = *c*}. In this case, the alternative hypothesis posited that clonal relationship induced a more homogeneous differentiation in terms of cell types, leading to a decreased number of cell types expected per families *T*(*D*). For this reason, by Monte Carlo approximation, we estimated the p-value for the left-tailed test as

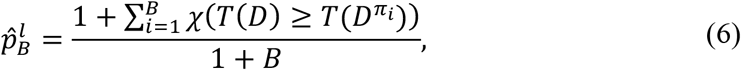

where *π*_1_,…,*π_B_* are sampled uniform and independent sampled elements from *Q*.

### Beta-binomial model for clonal concordance

For cells from the same ancestor, to quantify the correlation in decisions of cells from the same generation to continue to divide or cease dividing, we employed a stochastic mathematical model that was first described in [23]. The model is directly parameterized by the data, apart from one variable that encapsulates the correlation in decision-making that is fit to the data. In particular, with *n* being the maximum generation recorded, let *p_i_* ∈ [0,1] for *i* = 0,…,*n* denote the empirical proportion of cells that divide from generation *i* to the next, which is determined from the data as follows. Set *z_i_* the total number of cells recovered in generation *i*, the *p_i_* were estimated by

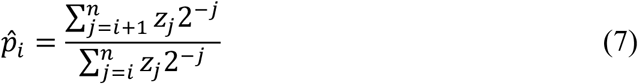

for *i* = 0,…,*n* − 1 and by 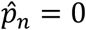.

In this model, given *k_i_* the number of cells from a particular family that reach generation *i*, the number of cells that continue on to divide to *i* + 1 follow a beta binomial distribution with parameters *k_i_, a_i_* = *p_i_*(1 − *ρ*)/*ρ*, and *b_i_* = (1 − *p_i_*)(1 − *ρ*)/*ρ*, namely *β*(*k_i_,a_i_,b_i_*), where *ρ* ∈ [0,1] is a free parameter. In particular, each family is generated recursively by setting *k*_0_ = 1 and defining *k*_*i*+1_ = 2*β*(*k_i_,a_i_,b_i_*). As in the experimental system, not all cells are recovered, but the proportion that are can be determined either by beads or well-volume recovered, on generating a family with the model, we accounted for sampling effect by subsampling each cell from a family with probability *r* = 0.71 independently of all other cells. The beta-binomial model interpolates between cells deciding to divide again independently of one another if *ρ* = 0, and when they are perfectly aligned, all making the same division decision, which occurs when *ρ* = 1. A value between 0 and 1 reflects the level of concordance within each family in division-progression decision-making, but, by construction, irrespective of the values of *p_i_*, that determines the population distribution among the generations. Defining the clonal range as the difference between maximum and minimum generations in which the cells from a family are recovered, the best-fit *ρ* was determined to be the value that maximized the likelihood of recapitulating the clonal range distribution in the data.

### Bootstrap confidence intervals

Confidence intervals at 95% level of Figures 2B-F and 3D were calculated via basic bootstrap [47]. To approximate the variability of the relevant statistic, 250,000 bootstrap datasets were created, each obtained by sampling with replacement the families that constituted the original data.

### Testing hypotheses on ancestral expressions and offspring progression or differentiation

Using the data underlying Figure 4D, we wished to challenge the null hypotheses that clonal progression is independent of ancestral expression levels (CD48, c-Kit, Sca-1). For notational purposes, the data were represented as a sequence 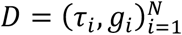 of *N* clones where the _i_^th^ family was identified by: expression level *τ_i_*, relative to one marker; maximum generation of its offspring *g_i_*. Given the set of maximum generations attained, *J* = {*g_i_,i* = 1,…,*N*], we partitioned the data into 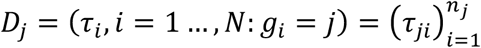 collections of size *n_j_*, for *j* ∈ *J*. Thus, we sought to test the null hypothesis that the variables *τ_ji_* are identically distributed, against the alternative hypothesis that, given *m_j_* the median of the distribution from which the elements of *D_j_* are drawn, for every *k, h* ∈ *J* such that *k* ≤ *h*, are either increasing

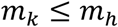

or decreasing

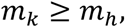

where at least one inequality must be strict. To this end, the statistic *T* of the data *D* was defined from the Jonckheere’s trend test [34], that is,

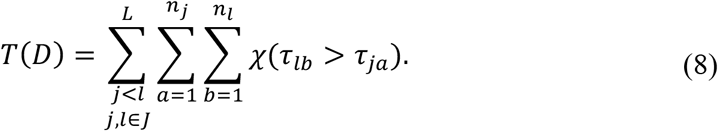

Under the null hypothesis that the variables *τ_ji_* are identically distributed, *D* is equally likely as a dataset 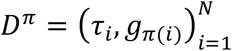 transformed by the action of any permutation *π* ∈ *Q* of the set of clonal labels {1,…,*N*}. As a consequence, using Monte Carlo approximation we estimated the p-value for the two-tailed test as

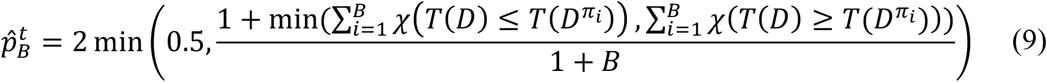

where *B* = 250,000 and *π*_1_,…,*π_B_* were uniformly and independently sampled from *Q*.

For the data underlying Figure 4E, we sought to challenge the null hypotheses that clonal differentiation to a certain cell type (SLAM-HSC, SLAM+ MEP, GMP, MEP, MPP) is independent of ancestral expression levels (CD48, c-Kit, Flt3, Sca-1). For these procedures, it sufficed to follow the same rationale as for the tests used for the data in Figure 4D, but for the dataset 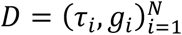 of *N* clones, where, for the *i*^th^ family, *τ_i_* is the expression level from one marker of its ancestor cell, while *g_i_* = 1 if its offspring was detected having at least one cell of the cell-type under consideration, *g_i_* = 0 otherwise. In particular, the testing statistic *T* was defined as in (8) and the subsequent p-value was estimated as in (9).

When multiple hypotheses were tested from the same data, the family-wise error rate was controlled using Holm-Bonferroni method [46]. As such, given the ordered p-values from *k* simultaneous tests 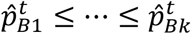, the *i*^th^ p-value was adjusted and recalculated as

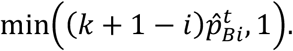

## Authorship

Study concept and design: L.P., K.D., T.T.; Experiment procedures and data acquisition: G.S., T.T., N.P.; data analysis: G.P., T.T, K.D. and L.P.; writing the manuscript: L.P, K.D, T.T and G.P.

## Acknowledgements

We would like to thank the members of Institut Curies Flow Facility for their help with setting up the flow cytometry experiments, the members of the Institut Curie Animalerie for their care for our experimental animals, Prof. Phil Hodgkin (Walter and Eliza Hall Institute of Medical Research) and Dr Julia Marchingo (University of Dundee) for their advice on setting up the experiment, and Stefania Pan and Emiliano Lancini (Université Paris 13) for their advice on optimization problems in graph theory. The present study was supported by an ATIP-Avenir grant from CNRS and Bettencourt-Schueller Foundation (to L.P.) and two grants from the *Labex CelTisPhyBio* (ANR-10-LBX-0038) and Idex Paris-Science-Lettres Program (ANR-10-IDEX-0001-02 PSL) (to L.P.).

## Supplementary Tables

**Supplementary Table 1.** Number of clones per progenitor and condition. Each entry of the table indicates the total number of clones or cells used to calculate the corresponding bar plot in Figure 2B-F, 2S and 4B,D. For Figure 4B,D, clones in different conditions were pooled. For the UMAP plots of Figure 3B-D and Figure 4A,C all clones were pooled, resulting in 1,592 cells.

| Condition (±IL-3/IL-6) | Progenitor | | | | | |
| --- | --- | --- | --- | --- | --- | --- |
|  | SLAM-HSC |  | ST-HSC |  | MPP |  |
|  | - | + | - | + | - | + |
| Total clones (Figure 2B,C and 4B,D) | 172 | 186 | 175 | 166 | 115 | 125 |
| Clones in generation 0 (Figure 2D) | 74 | 68 | 72 | 71 | 59 | 58 |
| Clones with 2 cells in generation 1 (Figure 2E) | 25 | 31 | 18 | 24 | 20 | 15 |
| Clones in recovered at 48 h time (Figure 2F) | 73 | 81 | 77 | 63 | 48 | 43 |
| Cells in generation 1 (Figure S2) | 76 | 90 | 60 | 65 | 63 | 46 |

**Supplementary Table 2.**
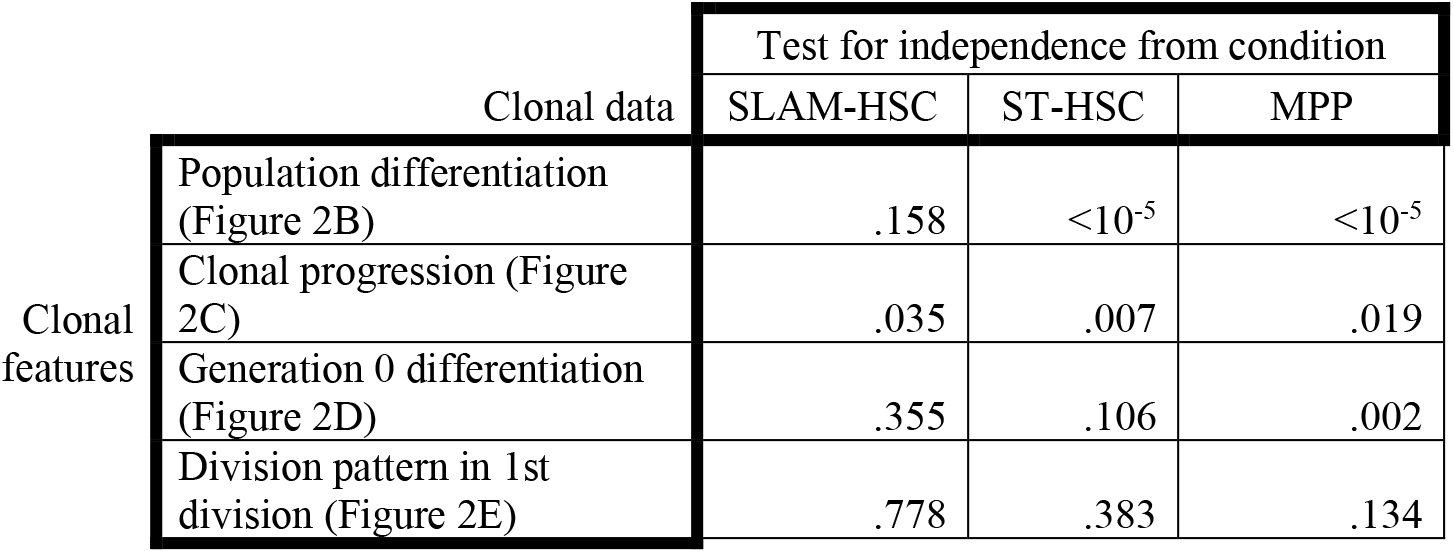
Significance values from permutation testing procedures. All the p-values from statistical tests carried out in Figure 2 are shown. Each value corresponds to a permutation test for independence of a clonal feature (one per row) with respect to either condition or progenitor structures, calculated on a given set of data (one per column).

**Supplementary Table 3.** Number of cells per generation from a given progenitor. Each entry of the table indicates the total number of cells used to calculate the corresponding proportion for the bar plot in Figure 3A.

| Progenitor | Generation |  |  |  |  |
| --- | --- | --- | --- | --- | --- |
|  | 0 | 1 | 2 | 3 | 4+ |
| SLAM-HSC | 142 | 166 | 151 | 161 | 29 |
| ST-HSC | 143 | 125 | 183 | 95 | 43 |
| MPP | 117 | 109 | 57 | 37 | 34 |

**Supplementary Table 4.** 95% confidence intervals of Spearman r and p-values of the correlation. All the values for Spearman correlation coefficient r, the 95% confidence intervals of r and the p value of the fit are shown for the correlations between fluorescence intensities of a given marker on each analyzed cell and the fluorescence intensity of the same marker on its’ ancestor cell at sort as shown in Figure 4E, reported for each ancestor cell separately.

| Marker | Progenitor |  |  |  |  |  |  |  |  |
| --- | --- | --- | --- | --- | --- | --- | --- | --- | --- |
|  | SLAM-HSC |  |  | ST-HSC |  |  | MPP |  |  |
|  | r | 95%CI | p | r | 95%CI | p | r | 95%CI | p |
| c-Kit | 0.007 | -0.073 - 0.086 | 0.864 | 0.207 | 0.126 - 0.285 | **<0.0001** | 0.333 | 0.236 - 0.425 | **<0.0001** |
| SLAM | 0.568 | 0.512 - 0.619 | **<0.0001** | 0.205 | 0.125 - 0.284 | **<0.0001** | 0.126 | 0.020 - 0.229 | **0.0164** |
| SCA1 | 0.603 | 0.550 - 0.651 | **<0.0001** | 0.519 | 0.456 - 0.577 | **<0.0001** | 0.567 | 0.490 - 0.635 | **<0.0001** |
| Flt3 | 0.110 | 0.031 - 0.188 | **0.0049** | 0.330 | 0.254 - 0.402 | **<0.0001** | 0.279 | 0.178 - 0.374 | **<0.0001** |
| CD48 | 0.650 | 0.592 - 0.701 | **<0.0001** | 0.473 | 0.390 - 0.548 | **<0.0001** | 0.305 | 0.188 - 0.413 | **<0.0001** |
| FSC | 0.061 | -0.019 - 0.139 | 0.1219 | 0.063 | -0.020 - 0.145 | 0.1269 | 0.074 | -0.0324 - 0.179 | 0.1603 |
| SSC | 0.003 | -0.076 - 0.082 | 0.9389 | -0.002 | -0.085 - 0.081 | 0.9555 | -0.105 | -0.209 - 0.001 | **0.0454** |

**Supplementary Table 5.** Significance values from the Jonckheere’s Trend test between expression levels at sort and offspring maximum generation. Each value corresponds to a p-value adjusted via Holm-Bonferroni correction (see Methods).

| SLAM-HSC | Ancestral marker expression | c-Kit | Sca-1 | CD48 |
| --- | --- | --- | --- | --- |
|  | - IL-3/IL-6, 24h | 0.1223 | 0.00496 | 0.000024 |
|  | + IL-3/IL-6, 24h | 0.6812 | 0.000216 | 0.000448 |
|  | - IL-3/IL-6, 48h | 0.001088 | 0.000768 | 0.5577 |
|  | + IL-3/IL-6, 48h | 0.000024 | 0.000016 | 0.00032 |

**Supplementary Table 6.** Significance values from the Jonckheere’s Trend test between expression levels at sort and presence of a given cell type among offspring. Each value corresponds to a p-value adjusted via Holm-Bonferroni correction (see Methods).

| Ancestor (time) | Ancestral marker | Condition and cell type |  |  |  |
| --- | --- | --- | --- | --- | --- |
|  |  | - IL-3/IL-6 |  | + IL-3/IL-6 |  |
|  |  | SLAM-HSC | SLAM+ MEP | SLAM-HSC | SLAM+ MEP |
| SLAM-HSC (24h) | CD48 | 0.07218 | 0.2305 | 0.000032 | 0.00228 |
|  | Sca-1 | 0.000048 | 0.00052 | 0.000048 | 0.00004 |
|  | c-Kit | 0.05708 | 0.10204 | 0.11032 | 0.3078 |
|  |  | SLAM-HSC | SLAM+ MEP | SLAM-HSC | SLAM+ MEP |
| SLAM-HSC (48h) | CD48 | 0.3711 | 0.3722 | 0.0153 | 0.04186 |
|  | Sca-1 | 0.000048 | 0.0094 | 0.000048 | 0.00016 |
|  | c-Kit | 0.03054 | 0.009696 | 0.004864 | 0.02358 |
|  |  | SLAM-HSC |  | SLAM-HSC |  |
| ST-HSC (24h) | CD48 | 0.8085 |  | 0.000304 |  |
|  | Sca-1 | 0.9432 |  | 0.01861 |  |
|  |  | SLAM-HSC | GMP | SLAM-HSC | GMP |
| ST-HSC (48h) | CD48 | 0.1047 | 0.000024 | 0.0255 | 0.000096 |
|  | Sca-1 | 0.13556 | 0.000144 | 0.03586 | 0.004176 |
|  |  | GMP | MEP | GMP |  |
| MPP (48h) | Sca-1 | 0.000048 | 0.6814 | 0.005376 |  |
|  | c-Kit | 0.09996 | 0.04688 | 0.013648 |  |
|  | Flt3 | 0.04772 | 0.9356 | 0.8337 |  |

**Figure S1.**
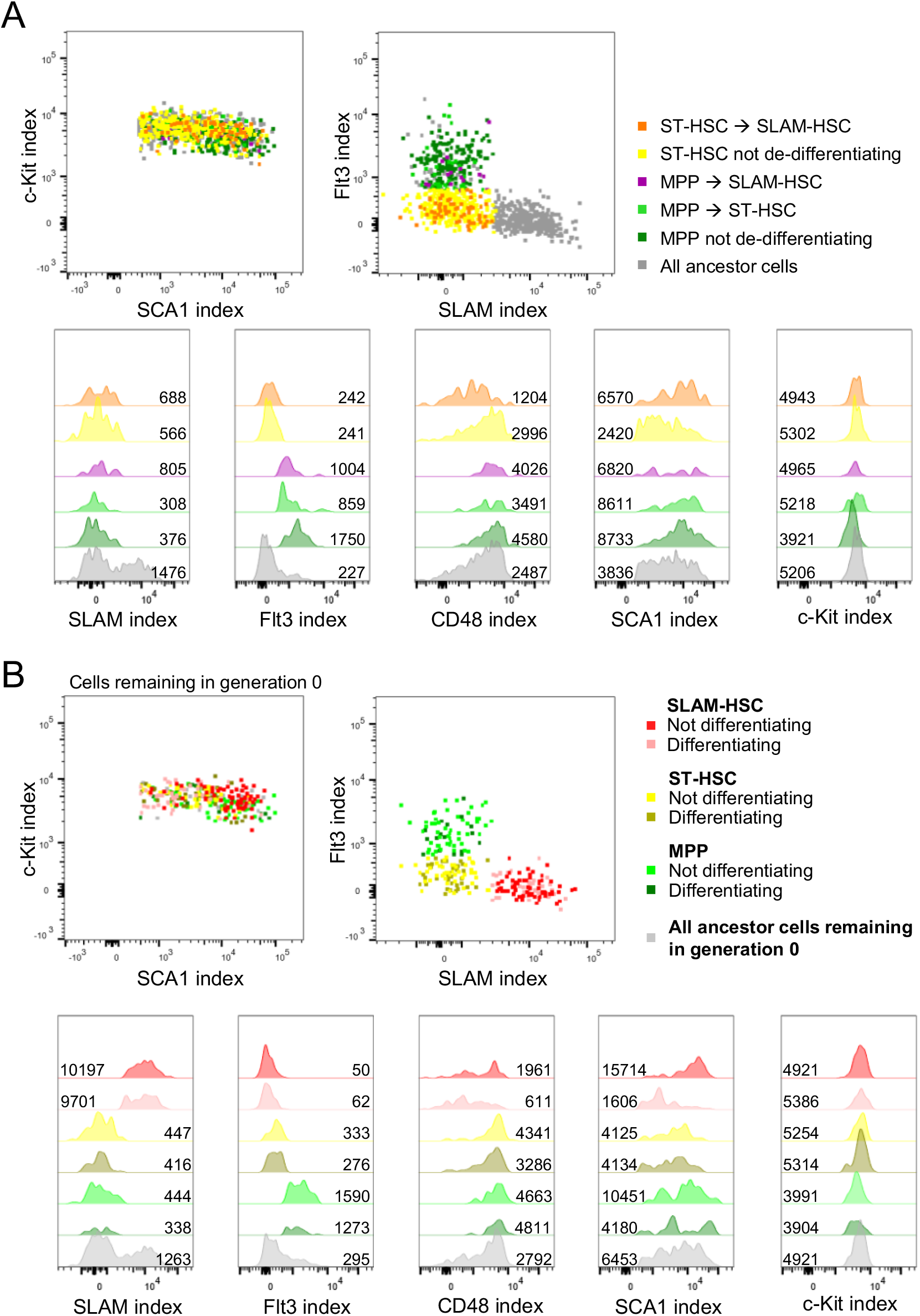
Index sorted fluorescence of differentiating, non-differentiating and dedifferentiating cells. (A) Fluorescence intensities at sort of dedifferentiating and non-dedifferentiating ancestors, both as 2D-plots and as histograms. Numbers next to histograms indicate median MFIs of the depicted population. (B) Fluorescence intensities at the sort of the differentiating and non-differentiating cells in generation zero from Figure 2D.

**Figure S2.**
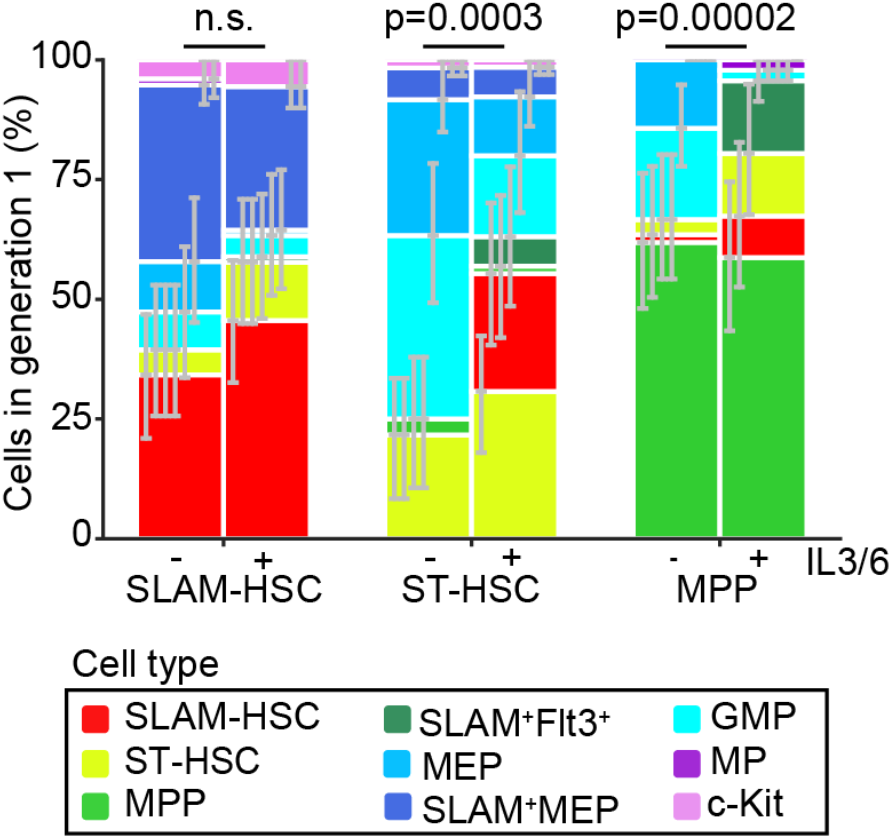
Proportions of recovered cell types for ancestors that have divided only once. Error bars indicate 95% confidence intervals calculated via basic bootstrap (Methods). Sample sizes for this panel can be found in Supplementary Table 1.

**Figure S3.**
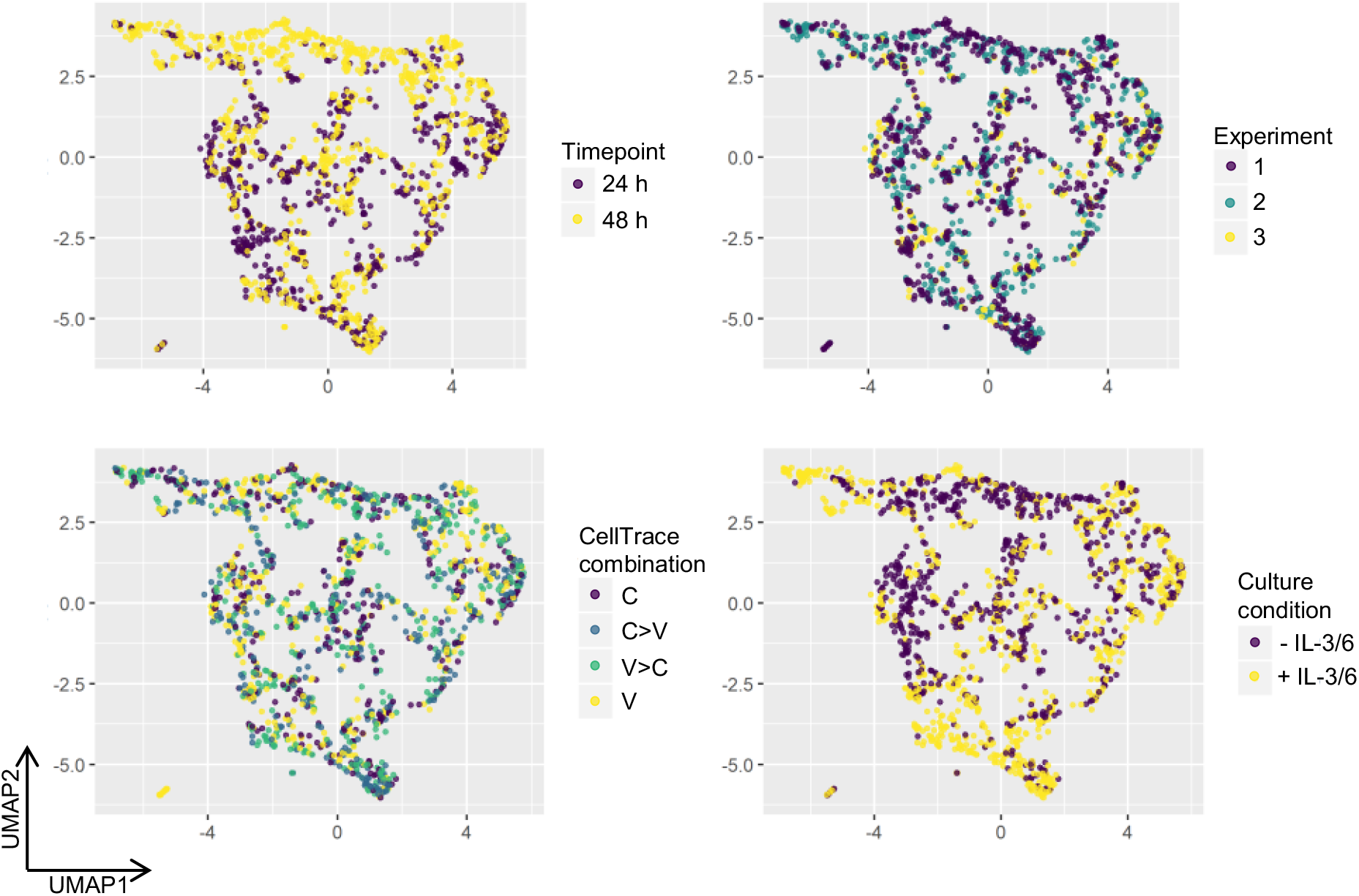
Additional information plotted onto the differentiation UMAP. Cells are color-coded by timepoint (top left), experiment (top right), CellTrace stain (bottom left) and culture condition (bottom right) and plotted on the UMAP from Figure 3. The experiments and CellTrace combinations were evenly distributed on the UMAP. Differences based on both the time-point and culture condition recapitulate what is described in Figures 2B and 3B. C indicates cells orgininating from ancestors stained stained with CFSE, V with CTV, C>V indicates staining with both 2.5 μM CFSE and 1.25 μM CTV and V>C indicates staining with both 1.25 μM CFSE and 2.5 μM CTV.

**Figure S4.**
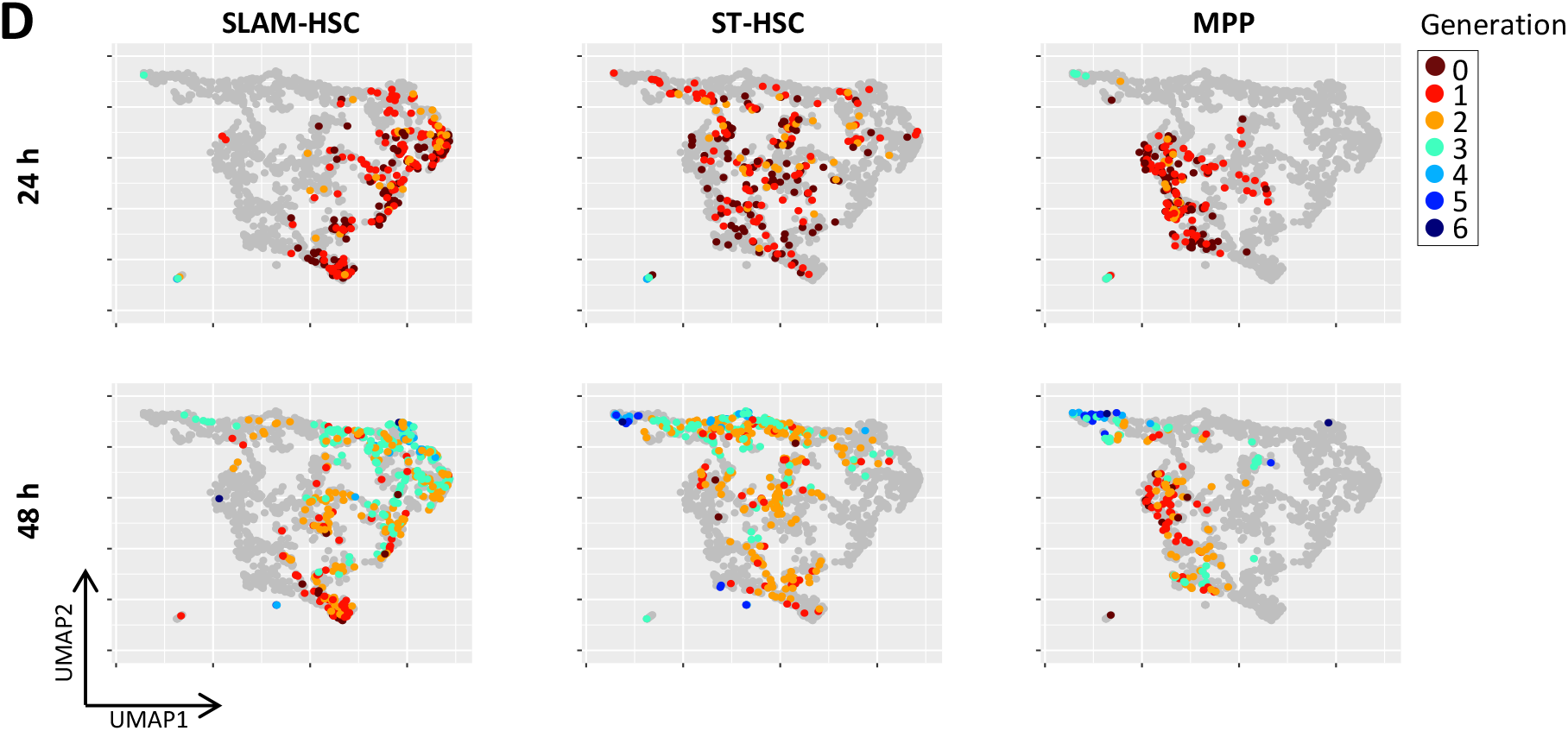
Generation numbers projected onto the UMAP, fractionated by ancestor type and time-point.

**Figure S5.**
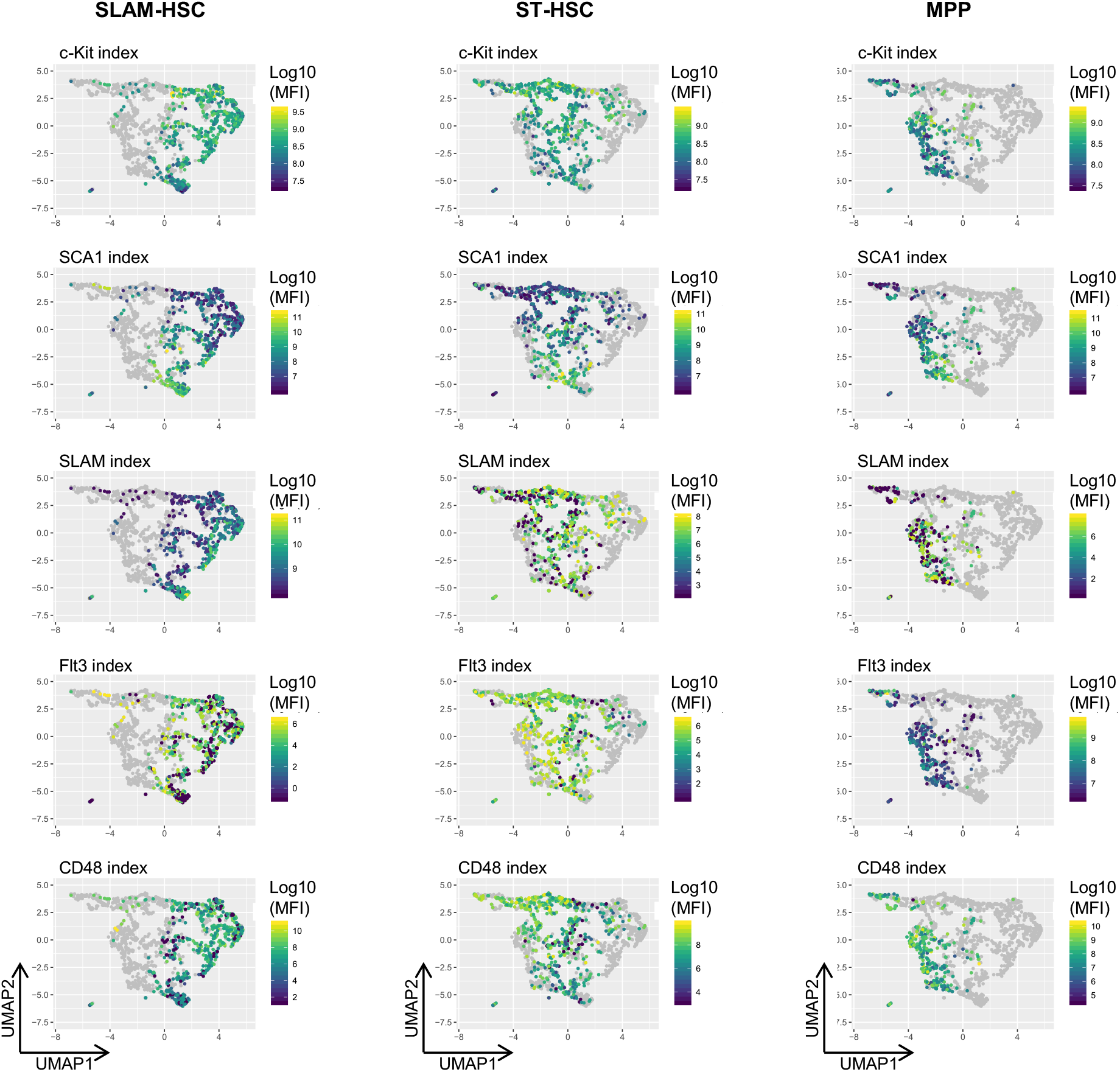
Per ancestor type fluorescence intensities during sort projected on the UMAP. As in Figure 4D, but fractionated by ancestor type.

**Figure S6.**
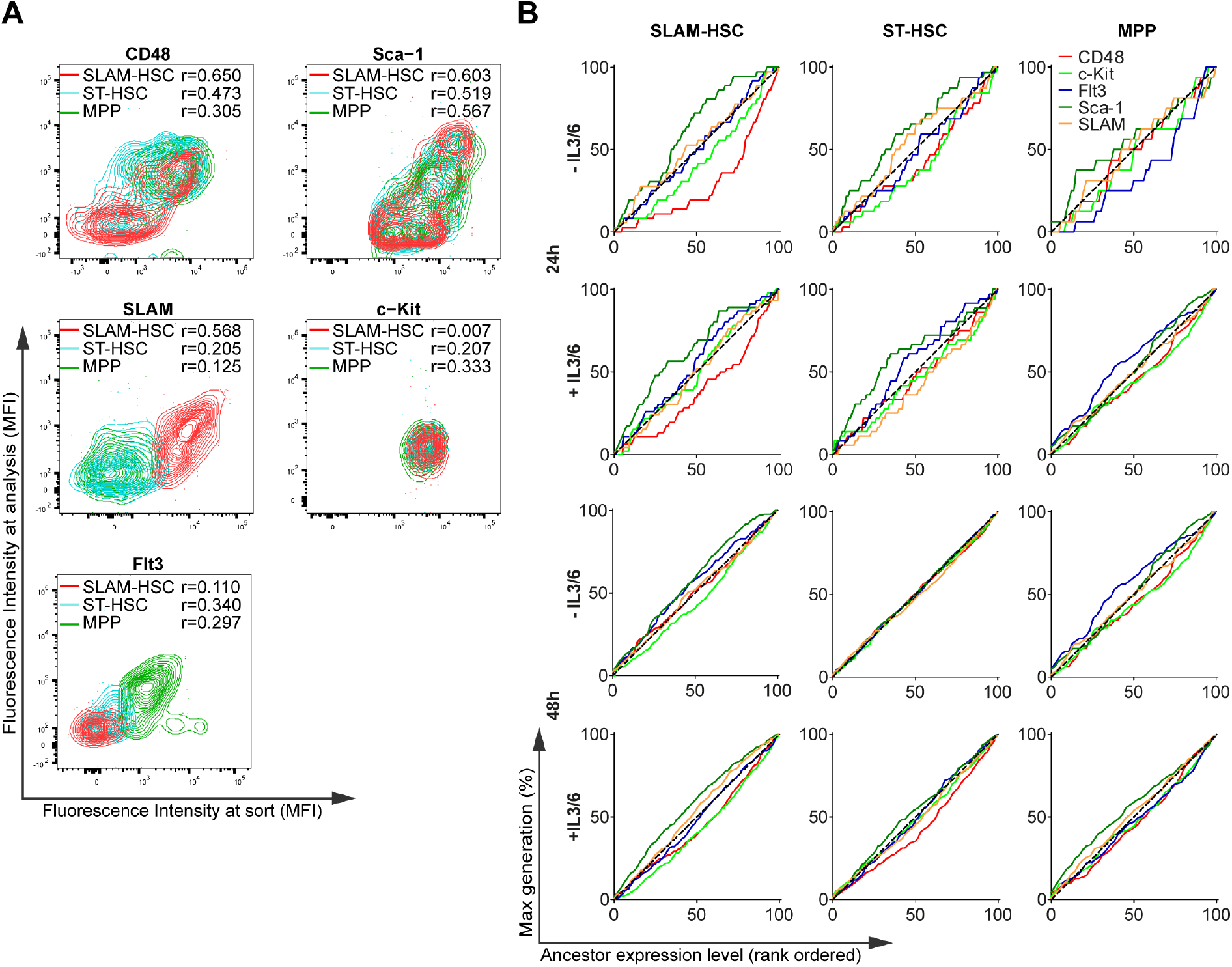
(A) Correlation between fluorescence intensities of the given marker on each analyzed cell and the fluorescence intensity of the same marker on its ancestor index sort. Spearman’s r is reported for index sort expression levels of each ancestor type. (B) The cumulative percentage of the maximum division of offspring from ancestor cells rank-ordered by their expression as in Figure 4D, but for each ancestor cell type and marker. Sample sizes for all panels can be found in Supplementary Table 1 and 95% CIs for the correlation r can be found in Supplementary Table 4.

**Figure S7.**
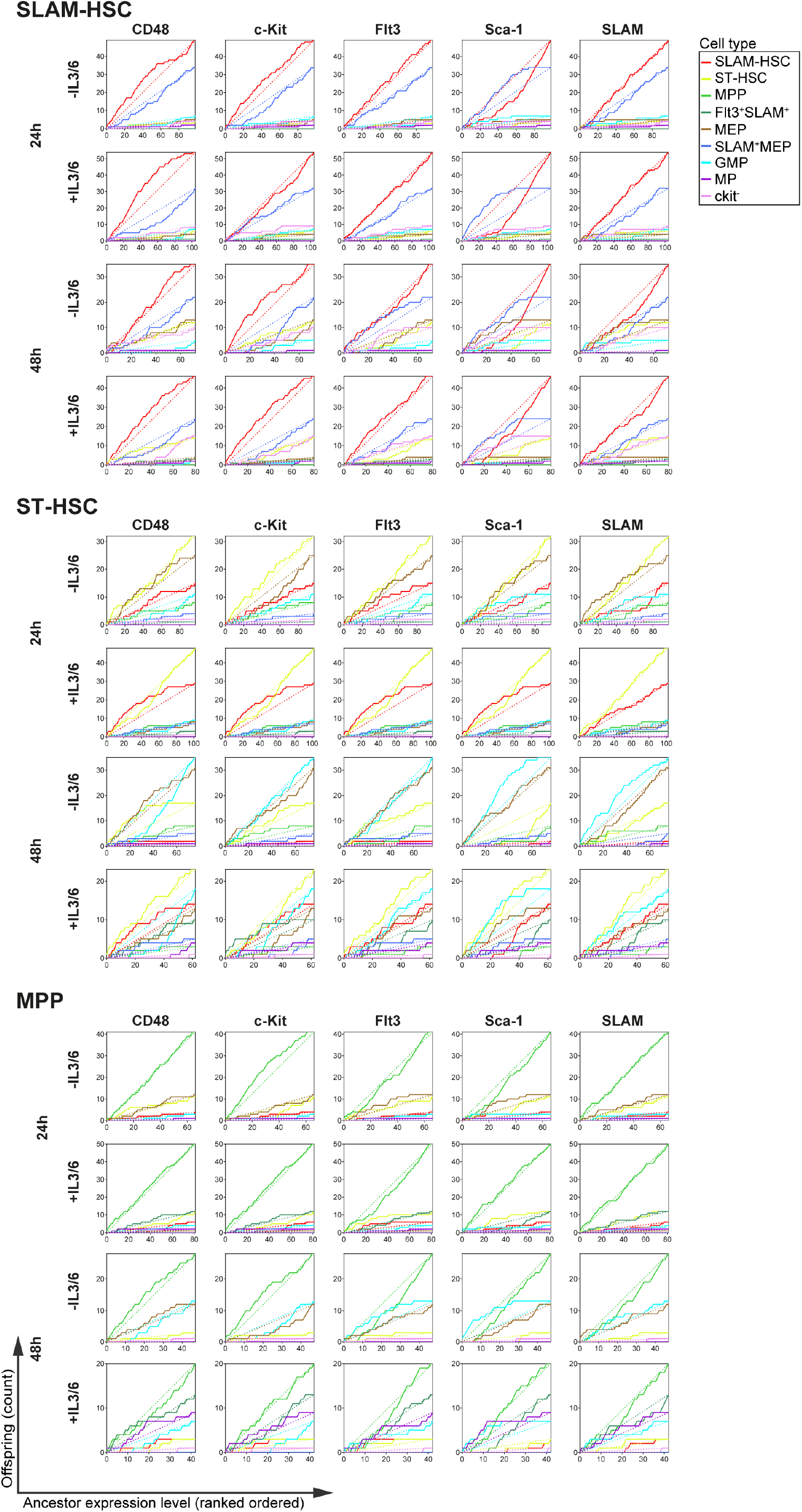
The cumulative sum of the offspring presenting a given cell type from ancestor cells rank-ordered by cell surface marker expression as in Figure 4E, without normalizing the offspring count, and plotted for each ancestor cell type, time point, culture condition and marker. Solid lines indicate observed sums, dashed lines indicate the diagonal for the corresponding observation.

